# Genetic signatures of evolution of the pluripotency gene regulating network across mammals

**DOI:** 10.1101/2020.04.16.043943

**Authors:** Yoshinori Endo, Ken-ichiro Kamei, Miho Inoue-Murayama

**Affiliations:** Wildlife Research Center, Kyoto University, 2-24 Tanaka-Sekiden-cho, Sakyo-ku, Kyoto, 606-8203, Japan; Institute for Integrated Cell-Material Sciences (WPI-iCeMS), Kyoto University, Yoshida-Ushinomiya-cho, Sakyo-ku, Kyoto, 606-8501, Japan; Wildlife Genome Collaborative Research Group, National Institute for Environmental Studies, 16-2 Onogawa, Tsukuba, Ibaraki, 305-8506, Japan

**Keywords:** Mammals, stem cells, pluripotency, gene network, evolution, natural selection

## Abstract

Mammalian pluripotent stem cells (PSCs) have distinct molecular and biological characteristics, but we lack a comprehensive understanding of regulatory network evolution in mammals. Here, we applied a comparative genetic analysis of 134 genes constituting the pluripotency gene regulatory network across 48 mammalian species covering all the major taxonomic groups. We found evolutionary conservation in JAK-STAT and PI3K-Akt signaling pathways, suggesting equivalent capabilities in self-renewal and proliferation of mammalian PSCs. On the other hand, we discovered rapid evolution of the downstream targets of the core regulatory circuit, providing insights into variations of characteristics. Our data indicate that the variations in the PSCs characteristics may be due to positive selections in the downstream targets of the core regulatory circuit. We further reveal that positively selected genes can be associated with species unique adaptation that is not dedicated to regulation of PSCs. These results provide important insight into the evolution of the pluripotency gene regulatory network underlying variations in characteristics of mammalian PSCs.

## Introduction

Pluripotent stem cells (PSCs) are undifferentiated cells that exhibit unlimited self-renewability and pluripotency, the potential to give rise to cells from all three embryonic germ layers. Pluripotent embryonic stem cells are isolated from the inner cell mass of developing pre-implantation mouse or human blastocysts (Evans & Kaufman 1981; Martin 1981; Thomson et al. 1998). Although PSCs can provide powerful resources for research and conservation of rare animal species, little equivalent derivations have been attempted with these species for ethical and technical reasons.

Recent advances in somatic cell reprogramming into induced PSCs (iPSCs) have broadened the opportunity to obtain PSCs from variety of mammals, including endangered species (Ben-Nun et al. 2011). The iPSC technology has been applied successfully to a wide range of taxonomic groups, including Carnivora (Shimada et al. 2009; Verma et al. 2012, 2013; Menzorov et al. 2015), Cetartiodactyla (Ezashi et al. 2009; Han et al. 2011; Liu et al. 2012), Chiroptera (Mo et al. 2014), Lagomorpha (Osteil et al. 2013), Metatheria (Weeratunga et al. 2018), Monotremata (Whitworth et al. 2019), Perissodactyla (Ben-Nun et al. 2011; Breton et al. 2013), Rodentia (Takahashi & Yamanaka 2006; Liao et al. 2009; Miyawaki et al. 2016; Lee et al. 2017), and Primates (Takahashi et al. 2007; Tomioka et al. 2010; Marchetto et al. 2013; Wunderlich et al. 2014; Ramaswamy et al. 2015). However, there continues to be discussion about species variation in the properties of the PSCs, such as pluripotent state, reprogramming efficiency, and optimal culture condition.

The characteristics of PSCs are controlled by a highly interconnected pluripotency gene regulatory network (PGRN) (Li & Belmonte 2017). Parts of the PGRN seem to be evolutionary conserved across mammals because iPSCs from different taxonomic groups have been derived using human genetic sequences of identical sets of transcription factors (*OCT4*, *SOX2*, *KLF4*, and *MYC*; collectively referred to as OSKM). On the other hand, different pluripotent states and configurations are observed across different species (Ezashi et al. 2016; Weinberger et al. 2016; Paterson et al. 2018), indicating diversity and uniqueness of the PGRN among species. However, our understandings of the PGRN are extensively limited to primates and rodents (Manor et al. 2015), rather than considering the evolutionary history of the mammalian PGRN across different taxonomic groups.

Comparative approaches can provide insights into the evolution of gene regulatory processes by examining the conservation and divergence of networks (Thompson et al. 2015). Evolutionary conservation and adaptations can be inferred by detecting purifying and positive selection (Nielsen et al. 2007). We have recently described genetic signatures for phenotypic adaptations in cetacean lipid metabolism (Endo et al. 2018), and numerous studies have revealed the effects of natural selection in the development of adaptive characteristics of animals and plants (Lenski 2017). Importantly, effects of changes on a regulatory process depends on the hierarchical position of the changes within the regulatory network (Erwin & Davidson 2009). Evolutionary pattern within the mammalian PGRN and how natural selection acts on the genes involved in the PGRN, however, are yet to be described.

In this study, we aimed to investigate whether any evolutionary pattern among mammalian taxa could be the result of selection of genes involved in the PGRN. To achieve this we conducted comparative selection analyses of the PGRN genes across mammals covering all the major taxonomic groups (**Figure 1**). (i) To assess the evolutionary conservation profiles of the mammalian PGRN, we estimated trends in the stringency of purifying selection and the evolutionary rate based on the ratio of substitution rates at non-synonymous and synonymous sites (d*N*/d*S*). (ii) To investigate the evolutionary force shaping the PGRN architecture, we compared the estimated conservation profiles among PGRN subcircuits. (iii) To detect phylogenetic inference of the variations in the PGRN, we identified genes under positive selection for lineages and investigated concordance in functional regions of the proteins. This paper presents the evolutionary history underlying the conservation and variations in mammalian pluripotency gene regulatory network, PGRN, and provides the genetic basis for the evolution of the PGRN in mammals.

**Fig. 1.**
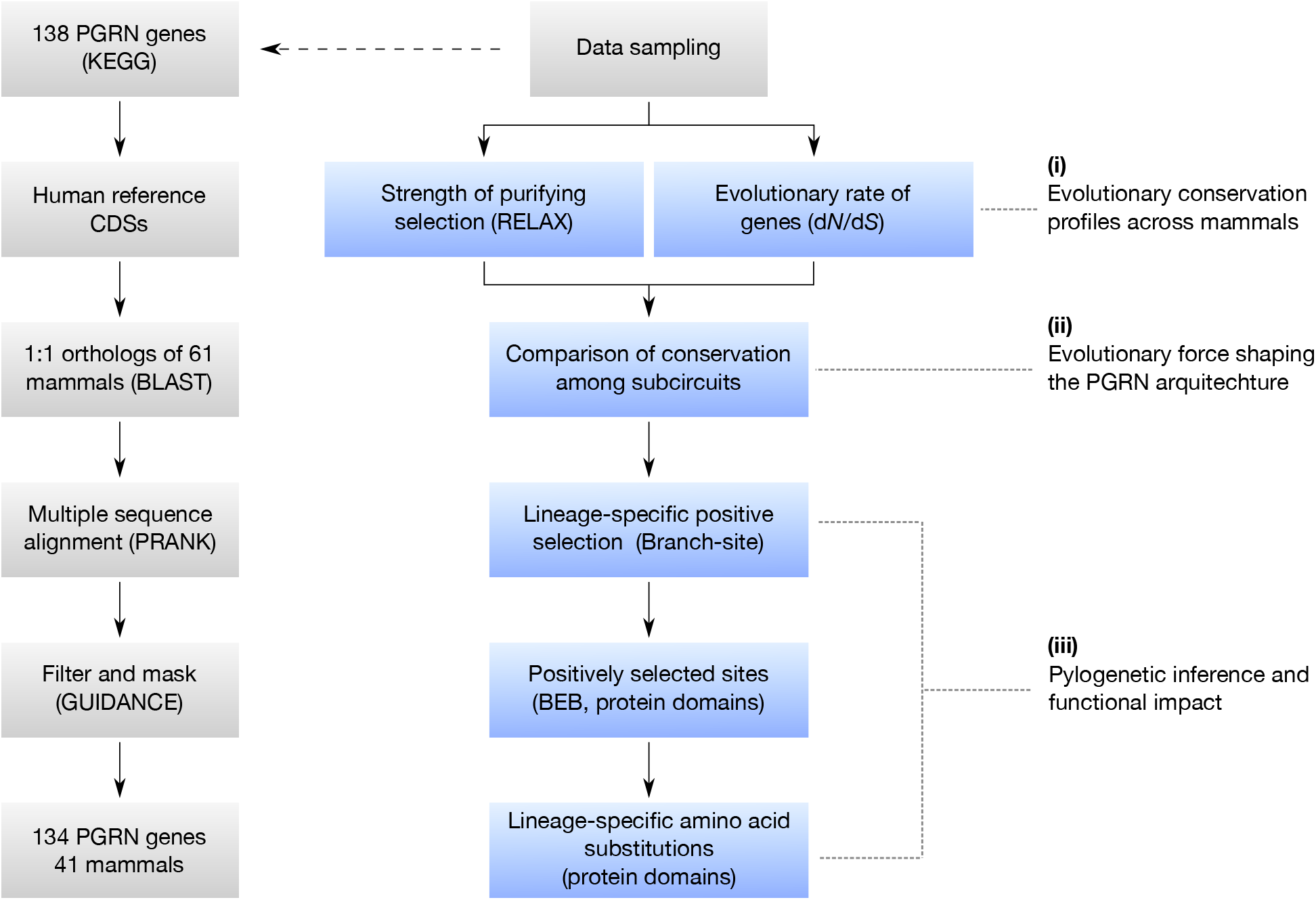
Schematic illustration of methods used to investigate the evolution of the PGRN across mammals.

## Methods

### Data sampling

Target genes were selected to include all 138 genes accomplishing “Signaling pathways regulating pluripotency of stem cells” (hsa04550) from Kyoto Encyclopedia of Genes and Genomes (KEGG) pathway database (Kanehisa et al. 2017). Human coding DNA sequences (CDS) of those genes were used as references and blasted against publicly available genomes of 61 mammals, covering all major taxonomic groups. *Gallus gallus*, *Xenopus tropicalis*, and *Danio rerio* were used as an outgroup. The set of genes were tested for orthologs using OMA stand alone v2.3.0 (Altenhoff et al. 2018). When a CDS was not inferred as an ortholog, another CDS was selected based on its annotation and tested with OMA again. Because the quality of some genomes might be poor, mammalian species with the same or lower number of orthologs than *Gallus gallus* were removed. Genes in which orthologs were found in less than 80% of the study species were also excluded from the further analyses. Orthologs were aligned with using PRANK v.140603 (Löytynoja & Goldman 2005) and the aligned sequences were further tested for their quality with GUIDANCE2 v2.02 (Sela et al. 2015) using the default settings. For the chicken genes, *POUV* (Gene ID: 427781, NP_001103648.1) and *NANOG* (Gene ID: 100272166, NP_001139614.1) were included in analyses even though they were not determined as orthologs by OMA because they are well studied and known as orthologs of human *POU5F1* and *NANOG* (Fuet & Pain 2017). A consensus tree was prepared based on commonly accepted phylogenies (Meredith et al. 2011; Perelman et al. 2011) and applied for later natural selection analyses. The gene ontology (GO) and the KEGG orthology (KO) were determined according to the Gene Ontology Annotation (GOA) and KEGG, respectively.

### Testing the stringency of natural selection

Intensity of natural selection for genes was tested by using RELAX (Wertheim et al. 2015) in HyPhy (Pond et al. 2005). RELAX is a general hypothesis testing framework that determines whether selective strength, distinguishing purifying or positive selection, was relaxed or intensified in the subsets of interest (Wertheim et al. 2015). RELAX estimates the selection intensity parameter, *k*, where *k* ≥ 1 indicates intensification and *k* < 1 indicates relaxation of natural selection where ω > 1 represents positive selection and ω < 1 represents purifying selection. Mammalian lineages were set as a test branch and non-mammalian vertebrates as a reference branch. Because RELAX requires reference branches, genes without non-mammalian orthologs were excluded. *P* values were corrected for False Discovery Rate (FDR) using the Benjamini-Hochberg method (Benjamini & Hochberg 1995) with a cutoff of 10%.

### Evolutionary rate of each gene

Evolutionary rate of each genes was estimated by using the model M0 in CodeML (PAMLv.4.8) (Yang 2007). The synonymous substitution rate (d*S*), the non-synonymous substitution rate (d*N*), and their ratio, d*N*/d*S* (also noted as ω) were calculated, where d*N*/d*S* < 1 indicates purifying selection, d*N*/d*S* = 1 indicates neutral, and d*N*/d*S* > 1 indicates positive selection. The d*N*/d*S* ratio was estimated for each gene using mammalian orthologs.

### Comparison of conservation patterns among subcircuits

The categories of subcircuits were determined according to KEGG descriptions in the “Pathway” category, including Core, JAK-STAT, MAPK, OSN activated, OSN suppressed, PI3K-Akt, TGFβ, and Wnt signaling pathways. As for the Core subcircuits, we included the genes in the core transcriptional network, *OCT4* (*POU5F1*), *SOX2*, and *NANOG* (OSN), and the genes which interact with the OSN inside of nucleus. When one gene was tagged in multiple subcircuits, it was grouped in all the related pathways. Because subcircuits contain unequal sample sizes, multiple paired comparisons for the means of d*N*/d*S* ratios were statistically tested using ANOVA followed by the Tukey-Kramer post-test with *P* value of 0.05.

### Lineage-specific positive selection

Positively selected genes (PSGs) were identified using the branch-site model (Zhang et al. 2005) implemented in CodeML (PAMLv.4.8) (Yang 2007). The modified model A as an alternative hypothesis was compared to the model A1 as the null. *P* values were evaluated under the likelihood ratio test (LRT) by comparing 2Δ*l* of the two models to the 1 : 1 mixture of 0 and χ^2^ distribution. Lineage-specific positive selection was tested by setting each contemporary species and ancestral branch as *a priori* specified target separately and corrected for FDR using the Benjamini-Hochberg method (Benjamini & Hochberg 1995) with a cutoff of 10%.

### Functional impact

Positively selected sites (PSSs) were determined using the Bayes empirical Bayes (BEB) implemented in the branch-site model (Yang et al. 2005) with 95% posterior probability. Lineage-specific amino acid substitutions were explored in PSGs identified in ancestral branches as indicative of significant changes in protein function (Tian et al. 2013). The lineage-specific amino acid substitutions were identified manually by looking at a conversion of residue that occurred and fixed in more than 90% of the descendant contemporary species Functional domain structures and regions were referred to UniPort and Pfam using human protein as a reference.

## Results

### Summary of sample data

To investigate the evolutionary pattern of the gene regulatory network, the PGRN, that maintains pluripotency and self-renewability of pluripotent stem cells, we performed comparative genetic analyses across mammalian species. We collected the protein coding sequences of PGRN related genes covering all the major taxonomic groups. A total of 61 mammalian species were used in this study including Primates, Rodentia, Cetartiodactyla, Chiroptera, Carnivora, Insectivora, Afrotheria, and Metatheria, with at least two species from each. For each species, we searched orthologs of the 138 genes assigned to functions associated with signaling pathways regulating pluripotency of stem cells. Our filtering strategy retained 134 gene sets from 48 mammalian species for later analyses (see **Methods**, **Supplementary Table S1**).

### Evolutionary conservation profiles across mammals

To assess evolutionary constraint and reduction of selective strength in the mammalian PGRN, we examined the strength of purifying selection with each gene. Of the 127 orthologous genes analyzed, we observed intensification of purifying selection in 53 genes (41.7%) as well as relaxation in 9 genes (7.1%) at *P* < 0.05 FDR 10% (**Figure 2**, **Supplementary Table S2**). Intensification of purifying selection was observed across the PGRN, while relaxation appeared to be limited in particular subcircuits.

**Fig. 2.**
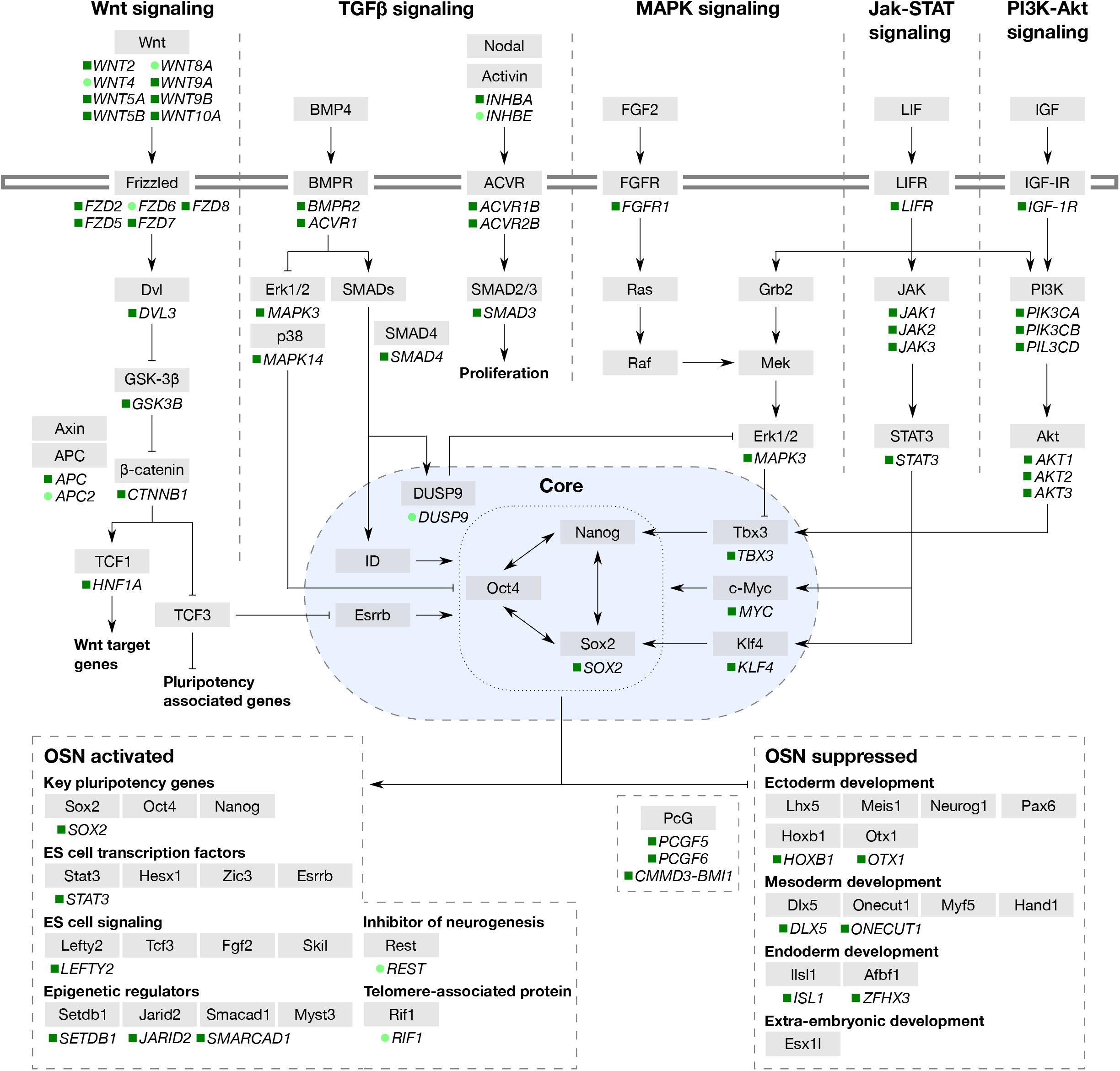
Trends in the stringency of purifying selection on the pluripotency regulating gene network (PGRN) genes. Schematic representation of the PGRN according to KEGG is shown. Only genes with significant shifts in the stringency of purifying selection at 10% FDR are indicated: *Green* rectangle, intensification; *lime* circle, relaxation. Dashed lines indicate the subcircuits of the PGRN. Horizontal double line indicates the cellular envelop. Blue oval represents the nucleus. The genes under each subcircuit are simplified, see **supplementary table S4** for details.

To gain insights into the main force that has shaped the evolution of mammalian PGRN, we estimated the evolutionary rates, d*N*/d*S*, of each gene. The estimated d*N*/d*S* values ranged from 0.0036 to 0.36. All the genes showed d*N*/d*S* values lower than 1 and 95.5% lower than 0.2 (**Figure 3A**, **Supplementary Table S3**).

**Fig. 3.**
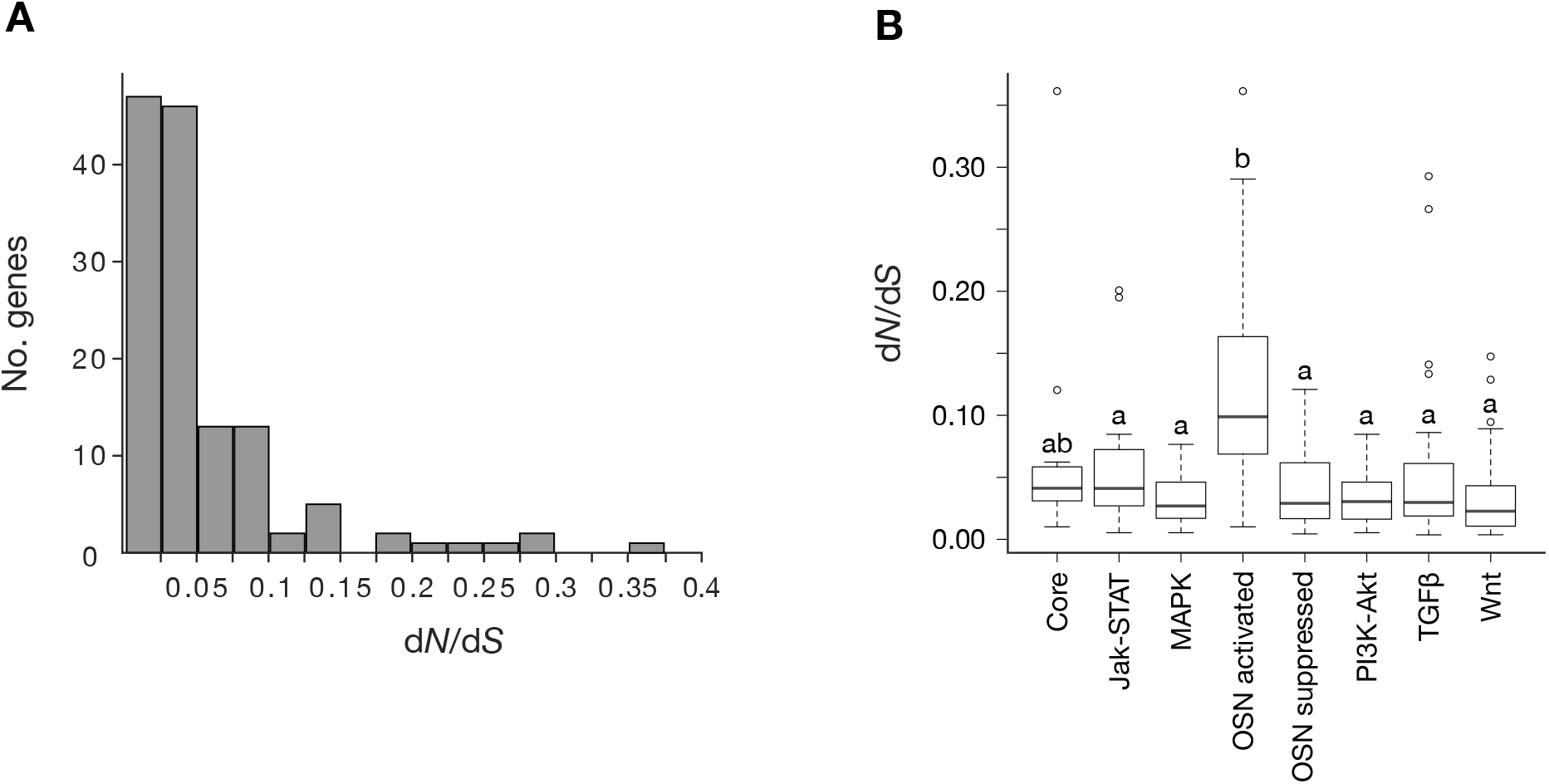
Conservation patterns of the pluripotency regulating gene network (PGRN). (*A*) Histogram of gene counts for d*N*/d*S* values showing distribution of evolutionary rates for the PGRN genes. (*B*) Box plots of the mean d*N*/d*S* values for the PGRN subcircuits. Multiple paired comparisons were tested with the Tukey-Kramer method, where subcircuits not assigned by the same latter are significantly different with *P* value of 0.05; error bars, 95% confidence interval. OSN represents *OCT4* (*POU5F1*), *SOX2*, and *NANOG*.

### Conservation patterns among the PGRN subcircuits

We then asked if there are differences in degree of conservation among subcircuits constituting the PGRN. In the purifying selection analysis above, we found relatively higher number of genes with intensified purifying selection in JAK-STAT, OSN suppressed, and PI3K-Akt signaling pathways (52.9%, 50%, and 44.8%, respectively), while more relaxed genes were found in OSN activated and Wnt signaling pathways (20% and 10%, respectively) (**Table 1**).

**Table 1.** Comparison of selective pressure among the subcircuits

| Subcircuits | No. gene | No. gene with significant shifts<br>in strength of purifying |  |
| --- | --- | --- | --- |
|  |  | Intensified | Relaxed |
| Core | 11 | 4 (36.4%) | 1 (9%) |
| JAK-STAT | 17 | 9 (52.9%) | 0 |
| MAPK | 26 | 8 (30.8%) | 1 (3.8%) |
| Core activated | 15 | 4 (26.7%) | 3 (20%) |
| Core suppressed | 18 | 9 (50%) | 0 |
| PI3K-Akt | 29 | 13 (44.8%) | 0 |
| TGFβ | 27 | 9 (33.3%) | 2 (7.4%) |
| Wnt | 40 | 16 (40%) | 4 (10%) |

We then examined the target genes based on the subcircuits that constitute the PGRN (**Supplementary Table S4**). Mean d*N*/d*S* values of each subcircuit were generally low; however, ANOVA indicated that evolutionary rates differ among subcircuits, and the following Tukey-Krammer showed that the OSN activated subcircuits, the downstream of the core transcriptional network, had significantly higher evolutionary rate compared to other subcircuits with *P* < 0.05 **(Figure 3B**, **Supplementary Table S5**).

### Lineage-specific positive selection

If genes in the OSN activated subdivision experience lower evolutionary constraint across mammals, some species may have developed unique characteristics through evolutionary changes in these genes. To test this, we performed positive selection analysis for the genes in the OSN activated subdivision on all the ancestral branches and contemporary species in the mammalian taxonomic tree containing our study species. Of the 15 OSN activated genes, we identified 8 PSGs in 4 ancestral branches and 22 contemporary species with 10% FDR correction (**Figure 4**, **Supplementary Table S6**). The four ancestral branches where PGSs were found include the common ancestor of Eutheria, Primates and the flying lemur, Megachiroptera, and Pan. The number of positive selection events was the highest in the gene *TCF3* (one ancestral branch and eight contemporary species). While three of the four tested epigenetic regulators (KO: 03036) have shown to be under intensified purifying selection across mammals, a relatively high number of positive selection events with *KAT6A*, *SETDB1*, and *JARID2* were observed in Primates (five events over nine total events in Primates and over 11 events with these genes across mammals).

**Fig. 4.**
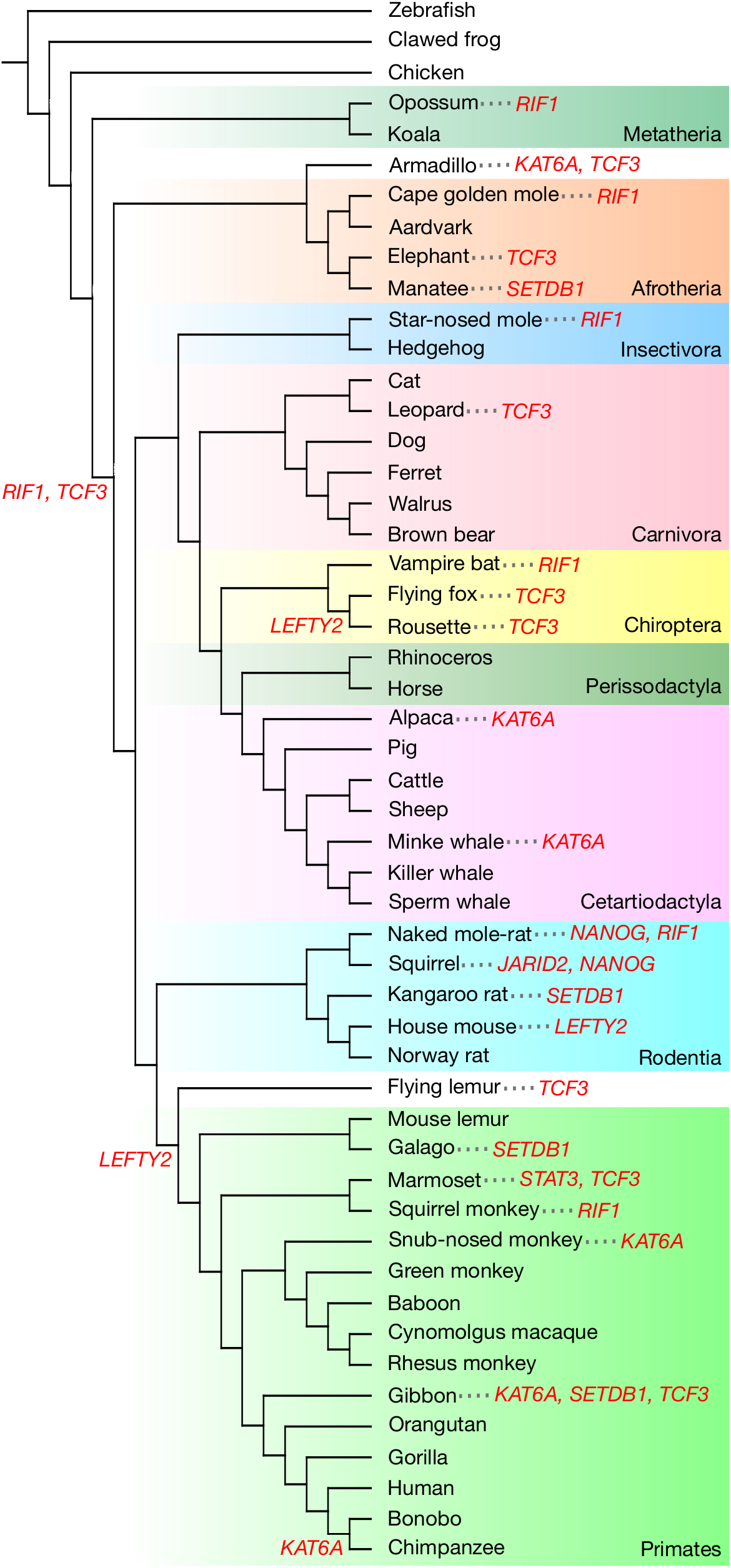
Mapping of positively selected genes identified in ancestral branches and contemporary species of mammals. Positively selected genes at 10% FDR (*red*) are indicated on a commonly accepted phylogeny of mammals. Colored boxes represents major taxonomic groups referred to the NCBI taxonomy.

### Functional impact of positively selected genes

To assess whether the identified PSGs have been functionally modified, we investigated the positively selected sites (PSSs) of the eight PSGs for each selected lineage (**Supplementary Table S7**). The positions of PSSs were compared to protein functional regions using humans as reference. We found sites under positive selection in functional regions of five proteins encoded by *KAT6A*, *LEFTY2*, *NANOG*, *SETDB1*, and *TCF3* (**Figure 5**). The majority of the PSSs were found outside of known functional regions. Although no significant PSSs were found in the PSGs at ancestral branches, we found lineage-specific amino acid substitutions in functional regions of all four PSGs, including *KAT6A*, *KEFTY2*, *RIF1*, and *TCF3*, in all the selected lineages (**Figure 6**, **Supplementary Table S8**).

**Fig. 5.**
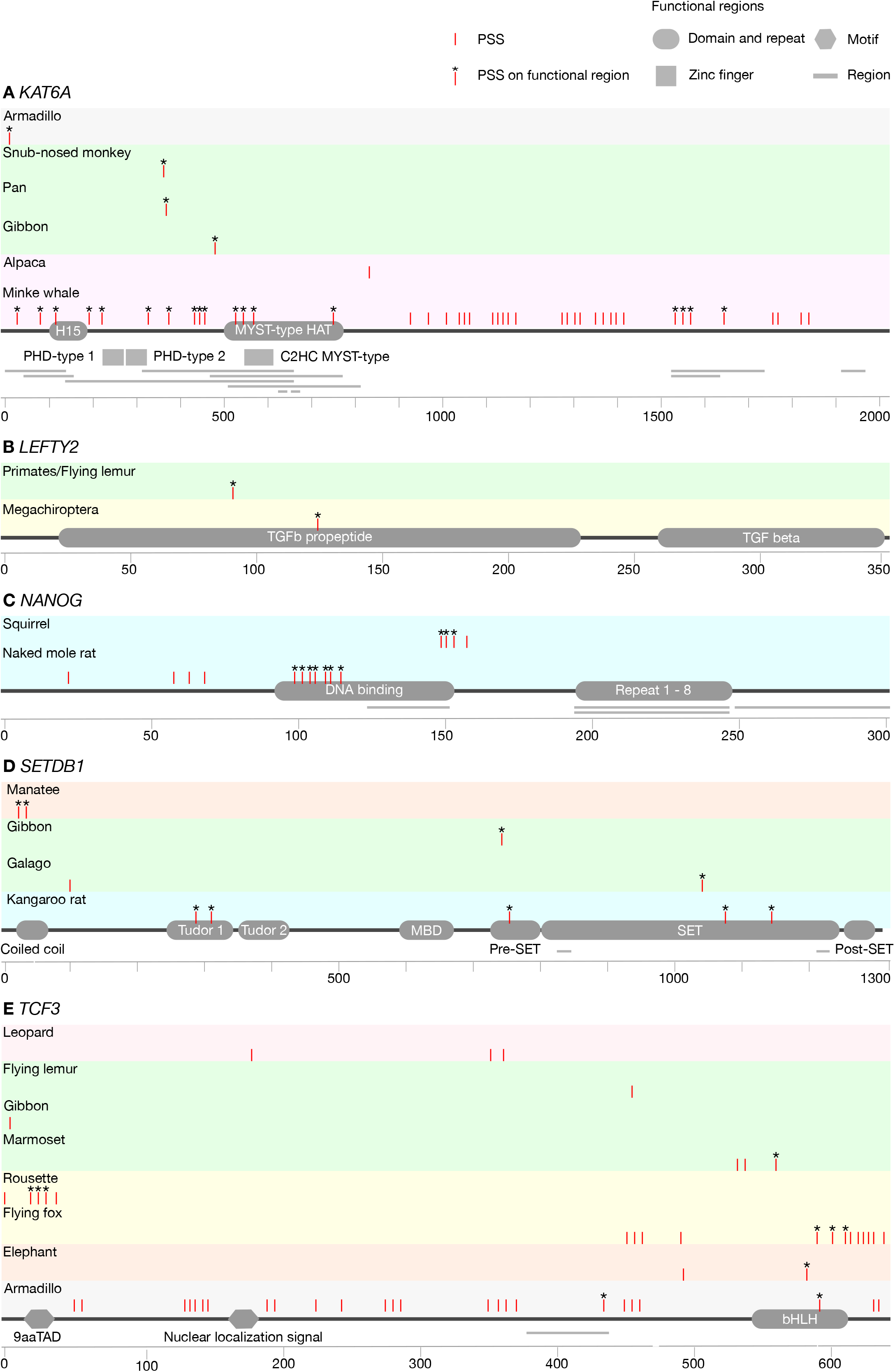
Positively selected sites (PSSs) of the OSN activated genes: (*A*) *KAT6A*, (*B*) *LEFTY2*, (*C*) *NANOG*, (*D*) *SETDB1*, and (*E*) *TCF3*. Positively selected lineages and their PSSs with 95% posterior probability are mapped on the coded protein with the arrangements of functionally important regions. Only genes with significant PSSs in at least one lineage and on at least one functional region are shown. The region represented with orange bar is a region of interest that cannot be described in other subsections according to UniPort. Colored boxes represent major taxonomic groups (Afrotheria (*orange*), Carnivora (*red*), Cetartiodactyla (*fuchsia*), Chiroptera (*yellow*), Primates/flying lemur (*lime*), Rodentia (*aqua*), and other (*grey*)).

**Fig. 6.**
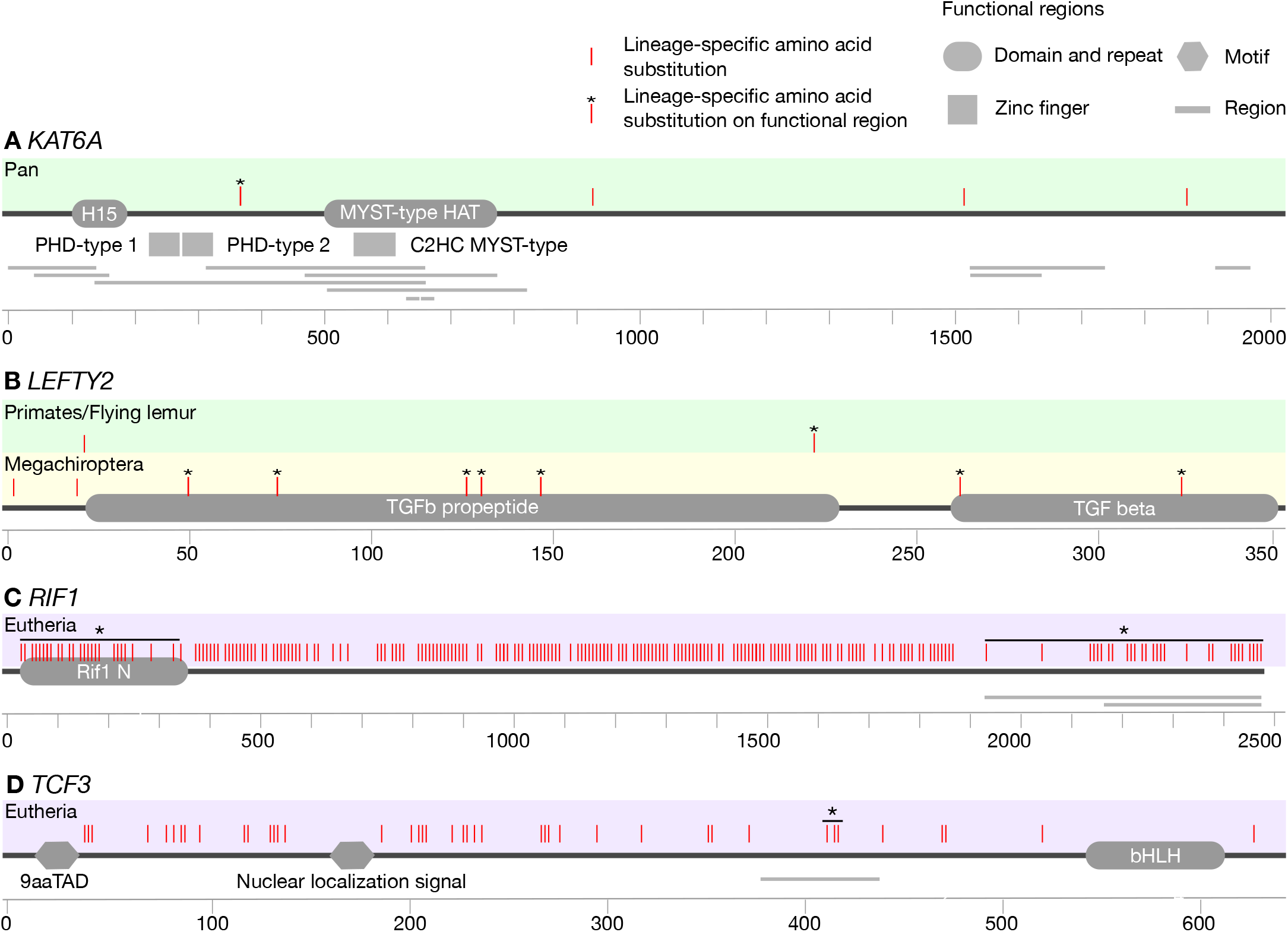
Lineage-specific amino acid substitutions in positively selected genes at ancestral branch: (*A*) *KAT6A*, (*B*) *LEFTY2*, (*C*) *RIF1*, and (*D*) *TCF3*. Positively selected lineages and their specific amino acid substitutions are mapped on the coded protein with the arrangements of functionally important regions. The region represented with orange bar is a region of interest that cannot be described in other subsections according to UniPort. Colored boxes represent major taxonomic groups (Eutheria (*purple*), Chiroptera (*yellow*), and Primates/flying lemur (*lime*)).

## Discussion

### Genetic conservation of the PGRN

We observed a high degree of sequence conservation among mammalian genes that constitutes the pluripotency gene regulatory network, PGRN. The prevalence of evolutionary conserved genes across the PGRN indicates overall comparability of the regulating mechanisms for pluripotency and self-renewability of mammalian PSCs. In context of the reprogramming somatic cells into induced pluripotent stem cells, although human sequences are effective (Ben-Nun et al. 2011), the efficient combination of reprogramming factors may differ among species (Tomioka et al. 2010; Verma et al. 2013; Mo et al. 2014; Weeratunga et al. 2018). Among the commonly used reprogramming factor, OSKM, we found genetic conservation of *SOX2*, *KLF4*, and *MYC* supporting the efficiency of these reprogramming factors in variety of species (Ezashi et al. 2016). On the other hand, our analyses did not indicate significant evidence for *POU5F1* and *NANOG*. Although *NANOG* is not necessary for reprogramming, it constitutes the core pluripotency transcription network of the PGRN (Li & Belmonte 2017). The binding regions of *POU5F1* and *NANOG* have been shown only ~5% similarity between humans and mice (Kunarso et al. 2010), and our findings may suggest that genetic sequences of these key regulators are poorly conserved across mammalian taxa. In addition to the reprogramming factors, the expression of other pluripotency-associated genes have been shown to enhance reprogramming efficiency (Takahashi & Yamanaka 2016). We detected significant evidence of purifying selection across mammals with *TBX3*, which improves reprogramming efficiency in mouse (Han et al. 2010). Considering suggested low sequence conservation in *POU5F1* and *NANOG*, our observations imply that *TBX3* might be a first candidate enhancer for derivation of mammalian iPSCs with low reprogramming efficiency.

The characteristics of mammalian PSCs may differ among species (Ezashi et al. 2016; Weinberger et al. 2016; Paterson et al. 2018) and reduction in selective constraint highlights evolutionary flexibility and innovation (Lahti et al. 2009; Moczek 2010; Hunt et al. 2011). For example, pluripotent state of PSCs can be described with multiple types such as naive or primed states, and the amenabilities of the pluripotent state *in vitro* differ between species (Boroviak et al. 2015). Although naive state PSCs are cultured with dual inhibition (2i) of MEK and GSK3 and leukemia inhibitory factor (LIF), human and mouse embryonic stem cells show distinct transcriptional responses to the 2i/LIF (Huang et al. 2014). We observed relaxation of purifying selection with genes involved in the molecular mechanisms that determine the pluripotent states (*APC2*, *DUSP9*, *INHBE*, *LEFTY2*, and *RIF1*). *RIF1* is involved in cellular response to LIF (GO: 1990830). *INHBE* and *LEFTY2* participate in the regulation of MAPK cascade (GO: 0043408), the target of MEK inhibitor, and the protein encoded by *DUSP9* is the essential regulator of MAPK (GO: 0000165, 0000187, 0000188). *APC2* is known to form a protein complex with the scaffold Axin and the kinases GSK3 and CK1 (Pronobis et al. 2015).

Interestingly, relaxation was observed in a number of oncogenes and tumor suppressor genes (*APC2*, *FZD6*, *INHBE*, *REST*, *RIF1*, *WNT4*, and *WNT8A*). Mammals lack correlation between body size or life span and cancer risk which is referred to as “Peto’s paradox” (Peto et al. 1975; Caulin & Maley 2011; Peto 2015). Our findings of the relaxation of selective strength for oncogenes and tumor suppressor genes may imply evolutional flexibility in mammalian cancer resistance. On the other hand, a high level of connectivity is proposed between positively selected genes (Vamathevan et al. 2008). Our findings of relaxation of purifying selection with a ligand *WNT4* and its receptor *FZD6* (Pronobis et al. 2015) may suggest a similar phenomena in relaxed genes.

Elements in fundamental processes, such as embryonic development, are expected to be conserved (Pennacchio et al. 2006). In agreement with this, we observed a high degree of evolutionary constraint among the PGRN genes with the skewed distribution of the evolutionary rates. A similar pattern has been reported for mammalian metabolic genes (Montanucci et al. 2018), but the evolutionary rates observed in the PGRN were considerably lower than those of metabolic genes, suggesting strong evolutionary conservation of the PGRN across mammals.

### Biased evolutionary conservation patterns among PGRN subcircuits

The patterns of genetic conservation among the PGRN subcircuits provide insight into the conservation and variations of the characteristics of mammalian PSCs. Multiple upstream signaling pathways serve to maintain the self-renewability and pluripotency of PSCs. We found relatively high degree of conservation in JAK-STAT, OSN-suppressed, and PI3K-Akt signaling pathways, suggesting conservation of fundamental biological characteristics and homeostasis in mammalian PSCs. JAK-STAT signaling pathway is stimulated by LIF and essential for self-renewal (Niwa et al. 1998). PI3K signaling pathway plays a crucial role for proliferation in mouse embryonic stem cells (Takahashi et al. 2005) and comparability of its function in mouse and primate ES cells has been reported (Watanabe et al. 2006) consistent with our observation. These findings may indicate that mammalian PSCs possess equivalent capabilities in self-renewal and proliferation. On the other hand, we observed the relatively higher number of relaxed genes in the Wnt signaling pathway. The downstream target of the Wnt signaling pathway, *ESRRB*, is necessary and sufficient to mediate self-renewal independently of JAK-STAT signaling pathway (Martello et al. 2012), that may suggest evolutionary flexibility in the parallel pathways supporting self-renewal. However, relaxed genes appear to be upstream of the Wnt signaling pathway (*WNT4A*, *WNT8A*, and *FZD6*), whereas the downstream catena of genes is under purifying selection (*GSK3B*, *CTNNB1*, and *HNF1A*), which illustrates a funnel-like distribution of the evolutionary pressures found in the network of metabolic genes (Montanucci et al. 2018), and subsequent influence on the PGRN is controversial.

The core transcriptional network genes, OSN, orchestrate a cascade of regulatory events involving an autoregulatory loop involving the other pluripotency regulators (Cole et al. 2008; Li & Belmonte 2017). We observed relatively high evolutionary rates with the genes activated by OSN. Because changes in the proximal targets of the master regulators could influence the subsequent circuitry (Erwin & Davidson 2009), our observations of relatively high evolutionary rates with the OSN activated subcircuit may reflect the variations of characteristics among mammalian PSCs. In addition, understanding mammalian diversity in the downstream targets of the core transcriptional network may help elucidate the mechanisms of reprogramming toward pluripotency and increase the reprogramming efficiency (Takahashi & Yamanaka 2016). Thus, we focused our later analyses on the genes in the OSN activated subcircuit for exploring variations in the mammalian PGRN.

### Development of lineage-specific PGRN

Evidence for positive selections on PGRN genes have implications for development of lineage-specific regulatory networks. We detected a relatively higher number of positive selections in contemporary species compared to in ancestral branches. This pattern contradicts that observed in a genome-wide scale study (Kosiol et al. 2008) indicating the unique development of the PGRN at a species level rather than at an order or a taxonomic group level. Species level development of the PGRN may explain the phylogenetic discrepancy of stem cell characteristics. Gene expression of the naked mole-rat iPSCs were more similar to that of human than to mouse iPSCs, despite their evolutionary relativeness (Lee et al. 2017). While the regulatory networks, and thus the gene expression patterns have been suggested to be context dependent (Hawkins et al. 2014; Weinberger et al. 2016), our sequence analyses do not experience this shortcoming.

Functional modifications in a transcription factor and its target provide further support for development of lineage-specific PGRN. *TCF3* encodes a member of the E protein family of helix-loop-helix transcription factors and plays a crucial role by binding to the component of the core transcription factors (Cole et al. 2008; Yi et al. 2008). We observed evidence of frequent episodes of positive selection with *TCF3*, the integral component of the core regulatory circuitry of PSCs, implying that alternation of the PGRN have occurred multiple times during mammalian evolution. For example, *ESRRB*, a regulator of self-renewal of PSCs (Martello et al. 2012), has been proposed to be recruited to the Eutheria PGRN after the divergence of marsupials and Eutheria because neither Tasmanian devil nor platypus iPSCs express *ESRRB* (Weeratunga et al. 2018; Whitworth et al. 2019). We found that the common ancestor of Eutheria has been positively selected with fixed lineage-specific amino acid substitutions with *TCF3*, which targets and controls the expression of *ESRRB* (Yi et al. 2008; Martello et al. 2012). We also report genetic evidence of positive selections with *TCF3* and its another target *LEFTY2* (Cole et al. 2008) in Megachiroptera. Positive selection has happened in the ancestral branch of Megachiroptera with *LEFTY2* and the two descendant species, the large flying fox (*Pteropus vampyrus*) and the rousette (*Rousettus aegyptiacus*) in *TCF3*, highlighting the evolutionary history of pre- and post-divergence of these taxa. Possible implications from this observation will be discussed in the next section. *TCF3* also regulates the expression of *NANOG* (Pereira et al. 2006; Cole et al. 2008). The reprogramming efficiency of *NANOG* varies among species; while overexpression of *NANOG* increases the reprogramming rate in felids (Verma et al. 2012, 2013), it does not increase the reprogramming rate in marmosets (Tomioka et al. 2010). We found that both the leopard and the marmoset have PSSs in their *TCF3*, but at different positions, which may reflect the variations of response to *NANOG*.

### Potential drivers of mammalian PGRN variation

Our findings of lineage-specific PSGs among the PGRN genes provide insight into the influence of species adaptation on the PRGN and the characteristics of mammalian stem cells. The architecture of developmental gene regulatory networks, such as the PGRN, is composed of diverse components. Certain subcircuits of regulatory networks are not dedicated to a particular biological process, but are also used for diverse functions that might have led to species adaptations. The naked mole-rat (*Heterocephalus glaber*) exhibits extraordinary longevity and cancer resistance (Buffenstein 2008; Delaney et al. 2013). The iPSCs derived from the naked mole-rat have shown to resist tumor formation through the expression of AFR (Miyawaki et al. 2016), in which the Arf/p53 pathway has a protective role from cancer and aging (Matheu et al. 2008). In agreement with this, we found *NANOG* and *RIF1* to be positively selected in this species. *RIF1* encodes a protein that regulates DNA replication and damage, interacting with 53BP1 (Kumar & Cheok 2014) which enhances p53 dependent transcriptional responses (Cuella-Martin et al. 2016). *NANOG* is not only one of the core transcriptional network genes, but also exhibits tumorigenic activity through interaction with p53 (Kim et al. 2016; Cuella-Martin et al. 2016). Our finding of PSSs on the DNA binding motifs of *NANOG* further suggests development of cancer resistance with p53 pathway in addition to AFR.

Alternatively, the evidence of positive selection on *RIF1* may indicate convergent evolution of species adaptation for perception. We identified that *RIF1* has been positively selected in the naked mole-rat, the Cape golden mole (*Chrysochloris asiatica*), the star-nosed mole (*Condylura cristata*), and the vampire bat (*Desmodus rotundus*). *RIF1* plays an important role in DNA replication and damage (Kumar & Cheok 2014), and these responses are important for neurogenesis, as represented in human diseases such as Meier-Grlin syndrome and Wolf-Hirschhorn syndrome (Kerzendorfer et al. 2013). Animals that live underground have developed unique sensory systems such as the somatosensory vibrissa-like body hairs on the body of the naked mole-rat (Crish et al. 2003), the middle ear structure of airborne and seismic stimuli in the Cape golden mole (Willi et al. 2006), and the “star” of the star-nosed mole (Gould et al. 1993). Among Microchiroptera that uses echolocation, the vampire bat has extremely sensitive neurons for noise that detect not only echolocation signals but also low frequency sounds, presumably for foraging prey in the dark (Schmidt et al. 1991).

Our results provide insights into additional possible development of cancer resistance in mammals. As discussed previously, two Megachiroptera species have been identified under positive selection with *TCF3*, which is also associated with cancer (Patel et al. 2015) and neuronal differentiation (Kuwahara et al. 2014). We observed multiple PSSs on the functional motif basic helix-loop-helix (bHLH) of *TCF3* in the large flying fox (*Pteropus vampyrus*), among the largest species of bat with a wingspan of up to 1.5m (Kunz & Jones 2000). Potential mechanisms of reducing cancer risk through response to DNA damage has been reported in elephants (Abegglen et al. 2015; Vazquez et al. 2018). Interestingly, we also observed PSSs on the bHLH of *TCF3* in elephants, implying the resolution to Peto’s paradox and the underlying convergent evolution between species that have developed a larger body size compered to their phylogenetic relatives. We identified PSSs on the transactivation domain, 9aaTAD in the rousette (*Rousettus aegyptiacus*), which has been reported to exhibit enhanced infection tolerance (Pavlovich et al. 2018). Whereas the previous study revealed genetic signatures of unique signaling in NK cell receptors (Pavlovich et al. 2018), E protein encoded by *TCF3* plays a critical role in B and T lymphocyte development (Engel et al. 2001; Seet et al. 2004), suggesting multiple strategies for antiviral defense in the rousette. Furthermore, we also detected positive selection with *TCF3* in the armadillo (*Dasypus novemcinctus*) with the highest number of PSSs. *TCF3* is also expressed in multipotent stem cells in skin, maintaining an undifferentiated state (Nguyen et al. 2006) and controlling cell fate (Merrill et al. 2001), pointing to a possible candidate for molecular evidence responsible for development of the outer shell.

The roles of transposable elements in genomic rearrangement, gene regulation, and epigenetics have been extensively studied to understand Primates evolution (Lee et al. 2016). Our findings of the relatively frequent positive selection with epigenetic regulators among Primates may reflect the various impacts of transposable elements on the primate genome. Among Primates, we observed that two epigenetic regulators were under positive selection in the gibbon (*Nomascus leucogenys*), whose chromatin interactions and epigenetic landscape has been remarkably conserved in spite of extensive genomic shuffling (Lazar et al. 2018). Interestingly, we also observed an event of positive selection at the common ancestor of Primates and the flying lemur with *LEFTY2*, with which DNA methylation plays a critical role during early embryogenesis in vertebrates (Wang et al. 2017), although the results need to be treated with caution.

Due to the habitat transition from terrestrial to aquatic environment, cetaceans have achieved a remarkable changes in their morphology (Uhen 2010). The minke whale (*Balaenoptera acutorostrata*) genome has provided support for genetic changes in *HOX* genes in this species (Yim et al. 2014), which have an important role in the body plan and embryonic development (Pearson et al. 2005). Consequently, we observed that the minke whale has been under positive selection with *KAT6A*, which regulates the expression of *HOX* gene (Voss et al. 2009), implying the morphological adaptation of the whale to the aquatic environment. However, our analyses did not indicate significant evidence of positive selection with *KAT6A* in other cetacean species. Because *KAT6A* is also associated with senescence and tumor growth (Baell et al. 2018), our findings with the minke whale may imply the adaptation of longevity and resistance to age related diseases as illustrated with the genome and transcriptomes of the bowhead whale (Keane et al. 2015).

Overall, our data indicated that the PGRN genes positively selected in some species are also involved in their unique adaptations, which may subsequently alter their regulatory function in PSCs. Further efforts, such as in vitro genetic modification and characteristics observation, are necessary to test the functional consequences of the genetic mutations discovered in this study.

## Conclusions

Our analyses illustrate the evolutionary patterns in the pluripotency gene regulatory network, PGRN, underlying the similarities and variations in characteristics among mammalian pluripotent stem cells, PSCs. This study is one of the first to compare the PGRN genes across major taxa. We showed the evolutionary conservation profiles of the mammalian PGRN and uncovered the evolutionary variable PGRN subcircuits. We identified phylogenetic inference of positive selection of genes involved in the PGRN which has enabled insights into development of lineage-specific PGRN and linkage between PGRN genes and species adaptation. These genes and the associated subcircuits will be plausible targets for future investigations exploring the mammalian PSCs.

## Authors’ contributions

Y. E. designed the original concept and scientific objectives of the project; performed data collection and data analyses; and wrote the manuscript. MI. M. and K. K. coordinated and participated in the design of the study, and helped draft the manuscript. All authors gave final approval for publication.

## Supplementary Material

Additional results supporting this article have been uploaded as part of the online electronic supplementary material.

## Acknowledgements

The authors thank Rob Ogden and Rebecca N. Johnson for constructive feedback on manuscript and English correction; Kristin Havercamp for valuable editorial input and English correction. This work was supported by JSPS KAKENHI Grant Number 17H03624 (MI-M) and 17H02083 (KK), and Kyoto University Supporting Program for Interaction-based Initiative Team Studies (SPIRITS) to MI-M.

## Literature Cited

Abegglen LM et al. 2015. Potential mechanisms for cancer resistance in elephants and comparative cellular response to dna damage in humans. JAMA. 314:1850–1860. doi: 10.1001/jama.2015.13134.

Altenhoff AM et al. 2018. The OMA orthology database in 2018: retrieving evolutionary relationships among all domains of life through richer web and programmatic interfaces. Nucleic Acids Res. 46:D477–D485. doi: 10.1093/nar/gkx1019.

Baell JB et al. 2018. Inhibitors of histone acetyltransferases KAT6A/B induce senescence and arrest tumour growth. Nature. 560:253–257. doi: 10.1038/s41586-018-0387-5.

Ben-Nun IF et al. 2011. Induced pluripotent stem cells from highly endangered species. Nat Methods. 8:829–831. doi: 10.1038/nmeth.1706.

Benjamini Y, Hochberg Y. 1995. Controlling the false discovery rate: a practical and powerful approach to multiple testing. J R Stat Soc Ser B. 57:289–300.

Boroviak T et al. 2015. Lineage-specific profiling delineates the emergence and progression of naive pluripotency in mammalian embryogenesis. Dev Cell. 35:366–382. doi: 10.1016/j.devcel.2015.10.011.

Breton A et al. 2013. Derivation and characterization of induced pluripotent stem cells from equine fibroblasts. Stem Cells Dev. 22:611–621. doi: 10.1089/scd.2012.0052.

Buffenstein R. 2008. Negligible senescence in the longest living rodent, the naked mole-rat: insights from a successfully aging species. J Comp Physiol B Biochem Syst Environ Physiol. 178:439–445. doi: 10.1007/s00360-007-0237-5.

Caulin AF, Maley CC. 2011. Peto’s paradox: evolution’s prescription for cancer prevention. Trends Ecol Evol. 26:175–182. doi: 10.1016/j.tree.2011.01.002.

Cole MF, Johnstone SE, Newman JJ, Kagey MH, Young RA. 2008. Tcf3 is an integral component of the core regulatory circuitry of embryonic stem cells. Genes Dev. 22:746–755. doi: 10.1101/gad.1642408.

Crish SD, Rice FL, Park TJ, Comer CM. 2003. Somatosensory organization and behavior in naked mole-rats I: Vibrissa-like body hairs comprise a sensory array that mediates orientation to tactile stimuli. Brain Behav Evol. 62:141–151. doi: 10.1159/000072723.

Cuella-Martin R et al. 2016. 53BP1 Integrates DNA Repair and p53-Dependent Cell Fate Decisions via Distinct Mechanisms. Mol Cell. 64:51–64. doi: 10.1016/j.molcel.2016.08.002.

Delaney MA, Nagy L, Kinsel MJ, Treuting PM. 2013. Spontaneous histologic lesions of the adult naked mole rat (*Heterocephalus glaberi*): A retrospective survey of lesions in a zoo population. Vet Pathol. 50:607–621. doi: 10.1177/0300985812471543.

Endo Y, Kamei K, Inoue-Murayama M. 2018. Genetic signatures of lipid metabolism evolution in Cetacea since the divergence from terrestrial ancestor. J Evol Biol. 31:1655–1665. doi: 10.1111/jeb.13361.

Engel I, Johns C, Bain G, Rivera RR, Murre C. 2001. Early thymocyte development is regulated by modulation of E2A protein activity. J Exp Med. 194:733–745. doi: 10.1084/jem.194.6.733.

Erwin DH, Davidson EH. 2009. The evolution of hierarchical gene regulatory networks. Nat Rev Genet. 10:141–148. doi: 10.1038/nrg2499.

Evans MJ, Kaufman MH. 1981. Establishment in culture of pluripotential cells from mouse embryos. Nature. 292:154–156. doi: 10.1038/292154a0.

Ezashi T et al. 2009. Derivation of induced pluripotent stem cells from pig somatic cells. Proc Natl Acad Sci. 106:10993–10998. doi: 10.1073/pnas.0905284106.

Ezashi T, Yuan Y, Roberts RM. 2016. Pluripotent stem cells from domesticated mammals. Annu Rev Anim Biosci. 4:223–253. doi: 10.1146/annurev-animal-021815-111202.

Fuet A, Pain B. 2017. Chicken induced pluripotent stem cells: establishment and characterization. In: Avian and Reptilian Developmental Biology: Methods and Protocols. Sheng, G, editor. Springer New York: New York, NY pp. 211–228. doi: 10.1007/978-1-4939-7216-6_14.

Gould E, McShea W, Grand T. 1993. Function of the star in the star-nosed mole, *Condylura cristata*. J Mammal. 74:108–116. doi: 10.2307/1381909.

Han J et al. 2010. Tbx3 improves the germ-line competency of induced pluripotent stem cells. Nature. 463:1096–1100. doi: 10.1038/nature08735.

Han X et al. 2011. Generation of induced pluripotent stem cells from bovine embryonic fibroblast cells. Cell Res. 21:1509–1512. doi: 10.1038/cr.2011.125.

Hawkins K, Joy S, McKay T. 2014. Cell signalling pathways underlying induced pluripotent stem cell reprogramming. World J Stem Cells. 6:620–628. doi: 10.4252/wjsc.v6.i5.620.

Huang K, Maruyama T, Fan G. 2014. The naive state of human pluripotent stem cells: A synthesis of stem cell and preimplantation embryo transcriptome analyses. Cell Stem Cell. 15:410–415. doi: 10.1016/j.stem.2014.09.014.

Hunt BG et al. 2011. Relaxed selection is a precursor to the evolution of phenotypic plasticity. Proc Natl Acad Sci U S A. 108:15936–15941. doi: 10.1073/pnas.1104825108.

Kanehisa M, Furumichi M, Tanabe M, Sato Y, Morishima K. 2017. KEGG: new perspectives on genomes, pathways, diseases and drugs. Nucleic Acids Res. 45:D353–D361. doi: 10.1093/nar/gkw1092.

Keane M et al. 2015. Insights into the evolution of longevity from the bowhead whale genome. Cell Rep. 10:112–122. doi: 10.1016/j.celrep.2014.12.008.

Kerzendorfer C, Colnaghi R, Abramowicz I, Carpenter G, O’Driscoll M. 2013. Meier-Gorlin syndrome and Wolf-Hirschhorn syndrome: Two developmental disorders highlighting the importance of efficient DNA replication for normal development and neurogenesis. DNA Repair (Amst). 12:637–644. doi: 10.1016/j.dnarep.2013.04.016.

Kim J, Liu Y, Qiu M, Xu Y. 2016. Pluripotency factor Nanog is tumorigenic by deregulating DNA damage response in somatic cells. Oncogene. 35:1334–1340. doi: 10.1038/onc.2015.205.

Kosiol C et al. 2008. Patterns of positive selection in six mammalian genomes. PLoS Genet. 4. doi: 10.1371/journal.pgen.1000144.

Kumar R, Cheok CF. 2014. RIF1: A novel regulatory factor for DNA replication and DNA damage response signaling. DNA Repair (Amst). 15:54–59. doi: 10.1016/j.dnarep.2013.12.004.

Kunarso G et al. 2010. Transposable elements have rewired the core regulatory network of human embryonic stem cells. Nat Genet. 42:631–634. doi: 10.1038/ng.600.

Kunz TH, Jones DP. 2000. Pteropus vampyrus. Mamm Species. 642:1–6. doi: 10.1644/0.642.1.

Kuwahara A et al. 2014. Tcf3 represses Wnt-β-catenin signaling and maintains neural stem cell population during neocortical development. PLoS One. 9:1–12. doi: 10.1371/journal.pone.0094408.

Lahti DC et al. 2009. Relaxed selection in the wild. Trends Ecol Evol. 24:487–496. doi: 10.1016/j.tree.2009.03.010.

Lazar NH et al. 2018. Epigenetic maintenance of topological domains in the highly rearranged gibbon genome. Genome Res. 28:983–997. doi: 10.1101/gr.233874.117.

Lee HE, Ayarpadikannan S, Kim HS. 2016. Role of transposable elements in genomic rearrangement, evolution, gene regulation and epigenetics in primates. Genes Genet Syst. 90:245–257. doi: 10.1266/ggs.15-00016.

Lee S-G et al. 2017. Naked mole rat induced pluripotent stem cells and their contribution to interspecific chimera. Stem Cell Reports. 9:1706–1720. doi: 10.1016/j.stemcr.2017.09.013.

Lenski RE. 2017. What is adaptation by natural selection? Perspectives of an experimental microbiologist. PLoS Genet. 13:1–12. doi: 10.1371/journal.pgen.1006668.

Li M, Belmonte JCI. 2017. Ground rules of the pluripotency gene regulatory network. Nat Rev Genet. 18:180–191. doi: 10.1038/nrg.2016.156.

Liao J et al. 2009. Generation of induced pluripotent stem cell lines from adult rat cells. Cell Stem Cell. 4:11–15. doi: 10.1016/j.stem.2008.11.013.

Liu J et al. 2012. Generation and characterization of reprogrammed sheep induced pluripotent stem cells. Theriogenology. 77:338–346. doi: 10.1016/j.theriogenology.2011.08.006.

Löytynoja A, Goldman N. 2005. An algorithm for progressive multiple alignment of sequences with insertions. Proc Natl Acad Sci U S A. 102:10557–10562. doi: 10.1073/pnas.0409137102.

Manor YS, Massarwa R, Hanna JH. 2015. Establishing the human naïve pluripotent state. Curr Opin Genet Dev. 34:35–45. doi: 10.1016/j.gde.2015.07.005.

Marchetto MCN et al. 2013. Differential L1 regulation in pluripotent stem cells of humans and apes. Nature. 503:525–529. doi: 10.1038/nature12686.

Martello G et al. 2012. Esrrb is a pivotal target of the Gsk3/Tcf3 axis regulating embryonic stem cell self-renewal. Cell Stem Cell. 11:491–504. doi: 10.1016/j.stem.2012.06.008.

Martin GR. 1981. Isolation of a pluripotent cell line from early mouse embryos cultured in medium conditioned by teratocarcinoma stem cells. Proc Natl Acad Sci U S A. 78:7634–7638. doi: 10.1073/pnas.78.12.7634.

Matheu A, Maraver A, Serrano M. 2008. The Arf/p53 pathway in cancer and aging. Cancer Res. 68:6031–6034. doi: 10.1158/0008-5472.CAN-07-6851.

Menzorov AG et al. 2015. Comparison of American mink embryonic stem and induced pluripotent stem cell transcriptomes. BMC Genomics. 16:S6. doi: 10.1186/1471-2164-16-s13-s6.

Meredith RW et al. 2011. Impacts of the Cretaceous Terrestrial Revolution and KPg extinction on mammal diversification. Science. 334:521–524. doi: 10.1126/science.1211028.

Merrill BJ, Gat U, DasGupta R, Fuchs E. 2001. Tcf3 and Lef1 regulate lineage differentiation of multipotent stem cells in skin. Genes Dev. 15:1688–1705. doi: 10.1101/gad.891401.

Miyawaki S et al. 2016. Tumour resistance in induced pluripotent stem cells derived from naked mole-rats. Nat Commun. 7:1–9. doi: 10.1038/ncomms11471.

Mo X, Li N, Wu S. 2014. Generation and characterization of bat-induced pluripotent stem cells. Theriogenology. 82:283–293. doi: 10.1016/j.theriogenology.2014.04.001.

Moczek AP. 2010. Phenotypic plasticity and diversity in insects. Philos Trans R Soc B Biol Sci. 365:593–603. doi: 10.1098/rstb.2009.0263.

Montanucci L et al. 2018. Influence of pathway topology and functional class on the molecular evolution of human metabolic genes. PLoS One. 13:1–17. doi: 10.1371/journal.pone.0208782.

Nguyen H, Rendl M, Fuchs E. 2006. Tcf3 governs stem cell features and represses cell fate determination in skin. Cell. 127:171–183. doi: 10.1016/j.cell.2006.07.036.

Nielsen R, Hellmann I, Hubisz M, Bustamante C, Clark AG. 2007. Recent and ongoing selection in the human genome. Nat Rev Genet. 8:857–868. doi: 10.1038/nrg2187.

Niwa H, Burdon T, Chambers I, Smith A. 1998. Self-renewal of pluripotent embryonic stem cells is mediated via activation of STAT3. Genes Dev. 12:2048–2060. doi: 10.1101/gad.12.13.2048.

Osteil P et al. 2013. Induced pluripotent stem cells derived from rabbits exhibit some characteristics of naive pluripotency. Biol Open. 2:613–628. doi: 10.1242/bio.20134242.

Patel D, Chinaranagari S, Chaudhary J. 2015. Basic helix loop helix (bHLH) transcription factor 3 (TCF3, E2A) is regulated by androgens in prostate cancer cells. Am J Cancer Res. 5:3407–3421.

Paterson YZ, Kafarnik C, Guest DJ. 2018. Characterization of companion animal pluripotent stem cells. Cytom Part A. 93:137–148. doi: 10.1002/cyto.a.23163.

Pavlovich SS et al. 2018. The Egyptian rousette genome reveals unexpected features of bat antiviral immunity. Cell. 173:1098–1110. doi: 10.1016/j.cell.2018.03.070.

Pearson JC, Lemons D, McGinnis W. 2005. Modulating Hox gene functions during animal body patterning. Nat Rev Genet. 6:893–904. doi: 10.1038/nrg1726.

Pennacchio LA et al. 2006. *In vivo* enhancer analysis of human conserved non-coding sequences. Nature. 444:499–502. doi: 10.1038/nature05295.

Pereira L, Yi F, Merrill BJ. 2006. Repression of Nanog gene transcription by Tcf3 limits embryonic stem cell self-renewal. Mol Cell Biol. 26:7479–7491. doi: 10.1128/mcb.00368-06.

Perelman P et al. 2011. A molecular phylogeny of living primates Brosius, J, editor. PLoS Genet. 7:e1001342. doi: 10.1371/journal.pgen.1001342.

Peto R. 2015. Quantitative implications of the approximate irrelevance of mammalian body size and lifespan to lifelong cancer risk. Philos Trans R Soc B Biol Sci. 370:20150198. doi: 10.1098/rstb.2015.0198.

Peto R, Roe FJ, Lee PN, Levy L, Clack J. 1975. Cancer and ageing in mice and men. Br J Cancer. 32:411–426. doi: 10.1038/bjc.1975.242.

Pond SLK, Frost SDW, Muse S V. 2005. HyPhy: hypothesis testing using phylogenies. Bioinformatics. 21:676–679. doi: 10.1093/bioinformatics/bti079.

Pronobis MI, Rusan NM, Peifer M. 2015. A novel GSK3-regulated APC: Axin interaction regulates Wnt signaling by driving a catalytic cycle of efficient βcatenin destruction. Elife. 4:1–31. doi: 10.7554/eLife.08022.

Ramaswamy K et al. 2015. Derivation of induced pluripotent stem cells from orangutan skin fibroblasts. BMC Res Notes. 8:577. doi: 10.1186/s13104-015-1567-0.

Schmidt U, Schlegel P, Schweizer H, Neuweiler G. 1991. Audition in vampire bats, *Desmodus rotundus*. J Comp Physiol A. 168:45–51. doi: 10.1007/BF00217102.

Seet CS, Brumbaugh RL, Kee BL. 2004. Early B cell factor promotes B lymphopoiesis with reduced interleukin 7 responsiveness in the absence of E2A. J Exp Med. 199:1689–1700. doi: 10.1084/jem.20032202.

Sela I, Ashkenazy H, Katoh K, Pupko T. 2015. GUIDANCE2: accurate detection of unreliable alignment regions accounting for the uncertainty of multiple parameters. Nucleic Acids Res. 43:W7–W14. doi: 10.1093/nar/gkv318.

Shimada H et al. 2009. Generation of canine induced pluripotent stem cells by retroviral transduction and chemical inhibitors. Mol Reprod Dev. 77:2–2. doi: 10.1002/mrd.21117.

Takahashi K et al. 2007. Induction of pluripotent stem cells from adult human fibroblasts by defined factors. Cell. 131:861–872. doi: 10.1016/j.cell.2007.11.019.

Takahashi K, Murakami M, Yamanaka S. 2005. Role of the phosphoinositide 3-kinase pathway in mouse embryonic stem (ES) cells. Biochem Soc Trans. 33:1522–1525. doi: 10.1042/BST20051522.

Takahashi K, Yamanaka S. 2016. A decade of transcription factor-mediated reprogramming to pluripotency. Nat Rev Mol Cell Biol. 17:183–193. doi: 10.1038/nrm.2016.8.

Takahashi K, Yamanaka S. 2006. Induction of pluripotent stem cells from mouse embryonic and adult fibroblast cultures by defined factors. Cell. 126:663–676. doi: 10.1016/j.cell.2006.07.024.

Thompson D, Regev A, Roy S. 2015. Comparative analysis of gene regulatory networks: From network reconstruction to evolution. Annu Rev Cell Dev Biol. 31:399–428. doi: 10.1146/annurev-cellbio-100913-012908.

Thomson JA et al. 1998. Embryonic stem cell lines derived from human blastocysts. Science. 282:1145–1147. doi: 10.1126/science.282.5391.1145.

Tian X et al. 2013. High-molecular-mass hyaluronan mediates the cancer resistance of the naked mole rat. Nature. 499:346–349. doi: 10.1038/nature12234.

Tomioka I et al. 2010. Generating induced pluripotent stem cells from common marmoset (*Callithrix jacchus*) fetal liver cells using defined factors, including Lin28. Genes to Cells. 15:959–969. doi: 10.1111/j.1365-2443.2010.01437.x.

Uhen MD. 2010. The origin(s) of whales. Annu Rev Earth Planet Sci. 38:189–219. doi: 10.1146/annurev-earth-040809-152453.

Vamathevan JJ et al. 2008. The role of positive selection in determining the molecular cause of species differences in disease. BMC Evol Biol. 8:1–14. doi: 10.1186/1471-2148-8-273.

Vazquez JM, Sulak M, Chigurupati S, Lynch VJ. 2018. A zombie *LIF* gene in elephants Is upregulated by TP53 to induce apoptosis in response to DNA damage. Cell Rep. 24:1765–1776. doi: 10.1016/j.celrep.2018.07.042.

Verma R et al. 2013. Nanog is an essential factor for induction of pluripotency in somatic cells from endangered felids. Biores Open Access. 2:72–76. doi: 10.1089/biores.2012.0297.

Verma R, Holland MK, Temple-Smith P, Verma PJ. 2012. Inducing pluripotency in somatic cells from the snow leopard (*Panthera uncia*), an endangered felid. Theriogenology. 77:220–228. doi: 10.1016/j.theriogenology.2011.09.022.

Voss AK, Collin C, Dixon MP, Thomas T. 2009. Moz and Retinoic Acid Coordinately Regulate H3K9 Acetylation, Hox Gene Expression, and Segment Identity. Dev Cell. 17:674–686. doi: 10.1016/j.devcel.2009.10.006.

Wang L et al. 2017. Epigenetic regulation of left–right asymmetry by DNA methylation. EMBO J. 36:2987–2997. doi: 10.15252/embj.201796580.

Watanabe S et al. 2006. Activation of Akt signaling is sufficient to maintain pluripotency in mouse and primate embryonic stem cells. Oncogene. 25:2697–2707. doi: 10.1038/sj.onc.1209307.

Weeratunga P, Shahsavari A, Ovchinnikov DA, Wolvetang EJ, Whitworth DJ. 2018. Induced pluripotent stem cells from a marsupial, the Tasmanian devil (*Sarcophilus harrisii*): insight into the evolution of mammalian pluripotency. Stem Cells Dev. 27:112–122. doi: 10.1089/scd.2017.0224.

Weinberger L, Ayyash M, Novershtern N, Hanna JH. 2016. Dynamic stem cell states: naive to primed pluripotency in rodents and humans. Nat Rev Mol Cell Biol. 17:155–169. doi: 10.1038/nrm.2015.28.

Wertheim JO, Murrell B, Smith MD, Kosakovsky Pond SL, Scheffler K. 2015. RELAX: Detecting relaxed selection in a phylogenetic framework. Mol Biol Evol. 32:820–832. doi: 10.1093/molbev/msu400.

Whitworth DJ et al. 2019. Platypus induced pluripotent stem cells: the unique pluripotency signature of a monotreme. Stem Cells Dev. 28:151–164. doi: 10.1089/scd.2018.0179.

Willi UB, Bronner GN, Narins PM. 2006. Ossicular differentiation of airborne and seismic stimuli in the Cape golden mole (*Chrysochloris asiatica*). J Comp Physiol A Neuroethol Sensory, Neural, Behav Physiol. 192:267–277. doi: 10.1007/s00359-005-0070-9.

Wunderlich S et al. 2014. Primate iPS cells as tools for evolutionary analyses. Stem Cell Res. 12:622–629. doi: 10.1016/j.scr.2014.02.001.

Yang Z. 2007. PAML 4: Phylogenetic analysis by maximum likelihood. Mol Biol Evol. 24:1586–1591. doi: 10.1093/molbev/msm088.

Yang Z, Wong WSW, Nielsen R. 2005. Bayes empirical Bayes inference of amino acid sites under positive selection. Mol Biol Evol. 22:1107–1118. doi: 10.1093/molbev/msi097.

Yi F, Pereira L, Merrill BJ. 2008. Tcf3 functions as a steady-state limiter of transcriptional programs of mouse embryonic stem cell self-renewal. Stem Cells. 26:1951–1960. doi: 10.1634/stemcells.2008-0229.

Yim H-S et al. 2014. Minke whale genome and aquatic adaptation in cetaceans. Nat Genet. 46:88–92. doi: 10.1038/ng.2835.

Zhang J, Nielsen R, Yang Z. 2005. Evaluation of an improved branch-site likelihood method for detecting positive selection at the molecular level. Mol Biol Evol. 22:2472–2479. doi: 10.1093/molbev/msi237.

